# A Perturbation Approach for Refining Boolean Models of Cell Cycle Regulation

**DOI:** 10.1101/2023.10.30.564745

**Authors:** Anand Banerjee, Asif Iqbal Rahaman, Alok Mehandale, Pavel Kraikivski

## Abstract

Considerable effort is required to build mathematical models of large protein regulatory networks. Utilizing computational algorithms that guide model development can significantly streamline the process and enhance the reliability of the resulting models. In this article we present a perturbation approach for developing data-centric Boolean models of cell cycle regulation. We assign a score to a network based on the steady states of the network-dynamics, and the dynamical trajectories corresponding to the initial conditions. Then, perturbation analysis is used to find new networks with lower scores, in which dynamical trajectories traverse through the correct cell cycle path with high frequency. We apply this method to refine Boolean models of cell cycle regulation in budding yeast and mammalian cells.

## INTRODUCTION

Cell cycle is the complex process through which a cell grows and divides into two genetically identical daughter cells. The progression of a cell through the cell cycle can be viewed as a trajectory through a multidimensional space of protein activation states which is controlled by an intricate network of biochemical reactions. The reaction network is fundamentally identical in all eukaryotes, and involves the activation/deactivation of different Cyclin-Cdk complexes at different stages of the cell cycle.

Most mathematical models of cell cycle are Ordinary Differential Equation (ODE) based [1-5]. While these models provide complete information about the time evolution of concentrations of different molecular species, they often require a large number of parameters in the form of reaction rate constants, whose values are often not known.

In studies of biological processes where such detailed temporal information is not necessary, Boolean modeling can be used to analyze the qualitative behavior of the system [6]. This network-based approach eliminates the need of kinetic parameters. The model consists of a network with nodes and edges. The nodes represent genes or proteins, and the edges signify reaction connecting them. Gene/protein activity is often represented by binary values 1 and 0 denoting active and inactive states respectively. These values are updated using logic-based rules. Boolean modeling has been used to study a wide range of biological phenomena including gene regulatory networks [7], signaling pathways [8], cell fate decisions [9], and cell cycle [10-13].

The update scheme in Boolean models has an important role in determining the dynamical behavior of the model [14]. In general, the update scheme can be defined as synchronous (also called deterministic) or asynchronous. In the synchronous update scheme, the values of all variables are updated simultaneously in one update step. The synchronous update is computationally easy to implement; however, it can produce spurious cycles in the dynamics. In contrast, in asynchronous update schemes, a randomly chosen variable is updated in a time step, resulting in stochastic representation of the dynamics. A drawback of the asynchronous scheme is that the trajectories can be very complicated and sometimes go through unrealistic pathways.

Among the Boolean models of cell cycle, Li et. al. [10] studied the cell cycle network in budding yeast using synchronous update scheme, and found that the primary steady state of the dynamics is stable to perturbations, and the globally attracting trajectory of the dynamics is consistent with sequence of protein states observed during cell cycle. Davidich et. al. [11], found similar results in the case of cell cycle networks in fission yeast. Faure et. al., used both synchronous and asynchronous update schemes to analyze mammalian cell cycle networks, and found that synchronous update resulted in two attractors: a steady state corresponding to G0 (with the inhibitors of cell cycle active), and a dynamical cycle corresponding to the cell cycle. Asynchronous update preserved the G0 steady state, but the cyclic attractor became very complex with many intertwined cycles.

Although these studies have laid the groundwork, they include only a small subset of genes/proteins involved in the cell cycle. Furthermore, the models were developed in a user-centric manner as opposed to data centric manner. To develop more comprehensive data-centric models, a systematic and automated approach is needed to construct the initial network, and then incorporate additional nodes into a growing network.

In this study, we aimed to develop a method for constructing Boolean models of cell cycle that transition through the correct sequence of the cell cycle stages with high frequency. To do so, we defined a network scoring method which depends on the steady states of the network as well as the dynamical trajectories corresponding to different initial conditions. All single and double perturbations of the networks were analyzed to search for new interactions that improve the network score and are also consistent with experimental data. We analyzed three cell cycle models shown in Fig.1: (a) Model A: the budding yeast cell cycle model [10], (b) Model B: the mammalian cell cycle regulation derived from Novak-Tyson model [1] and, (c) Model C: the mammalian cell cycle control derived from Gerard-Goldbeter model [2]. We found that many new interactions improve the score for each model. We searched for those interactions in databases to filter out the ones that are consistent with experimental data. Our future plan is to fully automate the process of comparing the interactions suggested by the perturbation algorithm with existing knowledge.

## RESULTS

### Budding yeast: Model A

We first analyzed the interaction network controlling the cell cycle in budding yeast, shown in Fig 1a, to test and optimize our perturbation-based approach. The network was taken from Li et.al. [10]. As mentioned previously, Li et.al., used synchronous update scheme and analyzed the robustness of the network by calculating the effect of single perturbations on the size of the largest attractor. We furthered that analysis by using the asynchronous update method, and a more comprehensive scoring scheme to quantify perturbations – one that incorporates the sizes of all the attractors, as well as the sequence of protein activation events.

**Figure 1.**
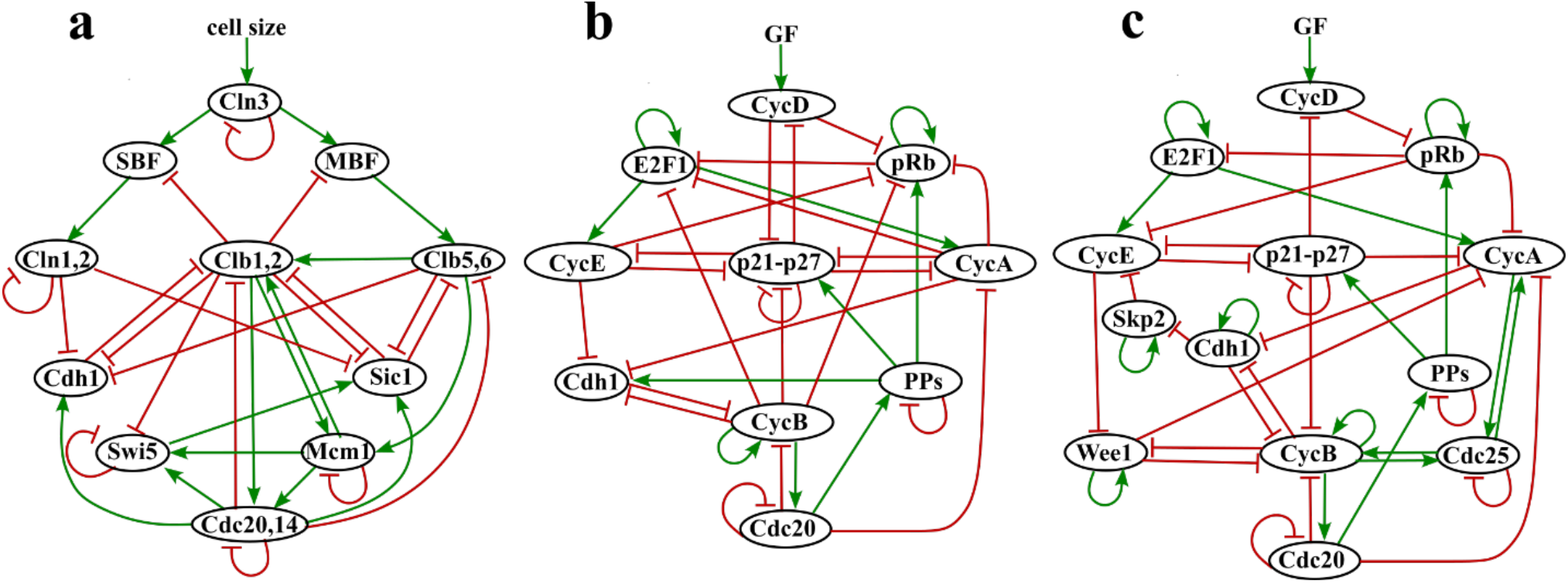
Three cell cycle models. (a) The budding yeast cell cycle regulation network derived from Li et. al. model [10]. (b), (c) Mammalian cell cycle regulation networks derived from Novak-Tyson and Gerard-Goldbeter models [1, 2], correspondingly. Oval shapes (network nodes) are proteins and lines between proteins (directed edges) are interactions that are of two types: activation represented by green arrow-headed lines and inhibition represented by red bar-headed lines.

**Figure 2:**
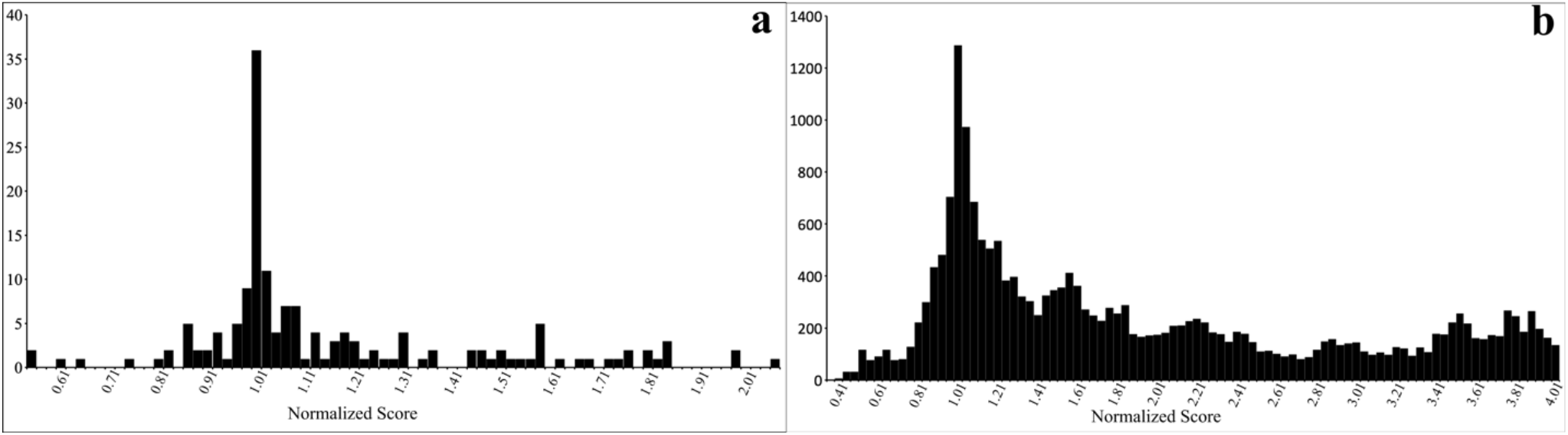
Histogram of normalized network scores after single perturbation (a) and double perturbation (b). The normalized scores were calculated by dividing the absolute network scores with score of the original network.

The network in Figure 1a has 11 nodes, resulting in 2^11^ = 2048 initial conditions. We found that all initial conditions eventually reach the 7 steady states listed in Table 1. These 7 states are the same as those found in Ref. Li et. al. [10], but because of the asynchronous updating rule used in our simulations, the size of the attractor corresponding to each steady state is slightly different in comparison. The largest attractor (in bold) corresponds to the G0 state.

**Table 1:** List of steady states and the corresponding number of initial conditions arriving at those steady states.

| Cln3 | MBF | SBF | Cln1,2 | Cdh1 | Swi5 | Cdc20,14 | Clb5,6 | Sic1 | Clb1,2 | Mcm1,SFF | Size of attractor |
| --- | --- | --- | --- | --- | --- | --- | --- | --- | --- | --- | --- |
| **0** | **0** | **0** | **0** | **1** | **0** | **0** | **0** | **1** | **0** | **0** | **1151** |
| 0 | 0 | 0 | 0 | 0 | 0 | 0 | 0 | 1 | 0 | 0 | 374 |
| 0 | 0 | 1 | 1 | 0 | 0 | 0 | 0 | 0 | 0 | 0 | 257 |
| 0 | 1 | 0 | 0 | 1 | 0 | 0 | 0 | 1 | 0 | 0 | 155 |
| 0 | 1 | 0 | 0 | 0 | 0 | 0 | 0 | 1 | 0 | 0 | 48 |
| 0 | 0 | 0 | 0 | 0 | 0 | 0 | 0 | 0 | 0 | 0 | 37 |
| 0 | 0 | 0 | 0 | 1 | 0 | 0 | 0 | 0 | 0 | 0 | 26 |

Interestingly, with asynchronous (or stochastic) updating, only 43% of the trajectories went through the correct sequence of protein activation. In 16% cases the trajectories were incorrect, and in 41% cases the trajectories ‘did-not-start’.

Next, we introduced perturbations to the network, with the aim of finding networks with a better score. The figure below shows the histogram of normalized scores after single and double perturbations to the network. We find that the peak of the distribution occurs at the score of the original graph, which is consistent with the Li et. al. However, interestingly, we also found perturbations which resulted in a lower score. The full list of network scores corresponding to single-edge and double-edge perturbations is given in the Supplementary Information File S1.

From the perturbations resulting in a score smaller than the original graph, the ones which are consistent with experimental data are listed in Table 2. For example, Skotheim et. al., observed that Cln1/2 dependent positive feedback promotes coherent SBF and MBF regulated gene expression [14]. Therefore, the addition of a positive influence of Cln1/2 on MBF can improve the consistency of the modeled cell cycle regulation with data. We also found supporting data for the other two interactions in Table 2. Whereas, it is well known that the transcription factor SBF controls CLN1/2 transcription, and the transcription factor MBF regulates CLB5/6 [15, 16], some data also suggest that in the absence of MBF, SBF is able to regulate expression of Clb5 [17]. Also, deletion analysis of upstream DNA sequences shows that Cln2 transcription can be induced by MBF transcription factor [18]. Furthermore, some known cell cycle models incorporate the positive regulation from SBF on Clb5,6 and MBF on Cln1,2 in addition to SBF→Cln1,2 and MBF→Clb5,6 [4, 19]. These additional interactions allowed the models to correctly characterize a vast amount of gene deletion mutant strains and better explain the regulation of cell cycle START transition.

**Table 2:** List of perturbations that are consistent with data and result in a network score smaller than the original network.

| Perturbation | Normalized Graph Score | steady state count | Largest attractor size | Correct | Incorrect | Did-not-start |
| --- | --- | --- | --- | --- | --- | --- |
| **OG Graph** | **1** | **7** | **1149** | **43%** | **16%** | **41%** |
| Single Perturbation |  |  |  |  |  |  |
| Cln1,2-to-MBF: 0 to 1 | 0.53 | 6 | 1321 | 56% | 20% | 24% |
| SBF-to-Clb5,6: 0 to 1 | 0.86 | 7 | 1277 | 46% | 29% | 25% |
| MBF-to-Cln1,2: 0 to 1 | 0.88 | 5 | 1297 | 53% | 22% | 25% |
| Clb5,6-to-Mcm1: 1 to 0 | 0.93 | 7 | 1162 | 58% | 0% | 42% |
| Double Perturbation |  |  |  |  |  |  |
| Cln1,2-to-MBF: 0 to 1<br>Clb5,6-to-Mcm1: 1 to 0 | 0.45 | 6 | 1332 | 78% | 0% | 22% |
| Cln1,2-to-MBF: 0 to 1<br>Cdh1 to Swi5: 0 to -1 | 0.53 | 6 | 1311 | 58% | 20% | 22% |
| MBF-to-Cln1,2: 0 to 1<br>SBF-to-Clb5,6: 0 to 1 | 0.74 | 5 | 1424 | 58% | 33% | 9% |

Interestingly, double perturbation Cln1,2-to-MBF: 0 to 1 and Clb5,6-to-Mcm1: 1 to 0 resulted in the largest drop in the network score. The combination of these perturbations improved the size of the largest attractor as well as the frequency with which the dynamics follows the correct trajectory. From the network it is clear that positive regulation of transcription factor MBF by Cln1,2 improves the chances of Clb5,6 activation, which marks the point-of-no-return for this model. This results in more trajectories going through the cell cycle. We also found that the steady state corresponding to the state {Cln3:0, MBF:0, SBF:1, Cln1,2:1, Cdh1:0, Swi5:0, Cdc2014:0, Clb5,6:0, Sic1: 0, Clb1,2:0, Mcm1,SFF: 0} is absent in the presence of this double perturbation.

### Mammalian cell: Model B

Next, we analyzed the ODE-based Tyson-Novak model [1] of cell cycle in mammalian cells. We constructed a Boolean model using the ODEs and introduced certain modifications to improve the cell cycle stability. The final network is shown in Fig. 1b. Our model has 10 nodes, resulting in 2^10^ = 1024 initial conditions. We found the largest attractor was the G0 state with a basin of attraction of size 283. The full list of steady states (101 in total) and the corresponding size of attractor are given in the Supplementary Information File S2.

Like previously, we used the perturbation analysis to find networks with a lower score. The distributions of scores after single-edge and double-edge perturbations are shown in Fig 3. The distribution again peaks at the original graph score, but in this case, we find more perturbations resulting in a lower graph score. The full list of network scores corresponding to single-edge and double-edge perturbations is given in the Supplementary Information File S2.

**Figure 3.**
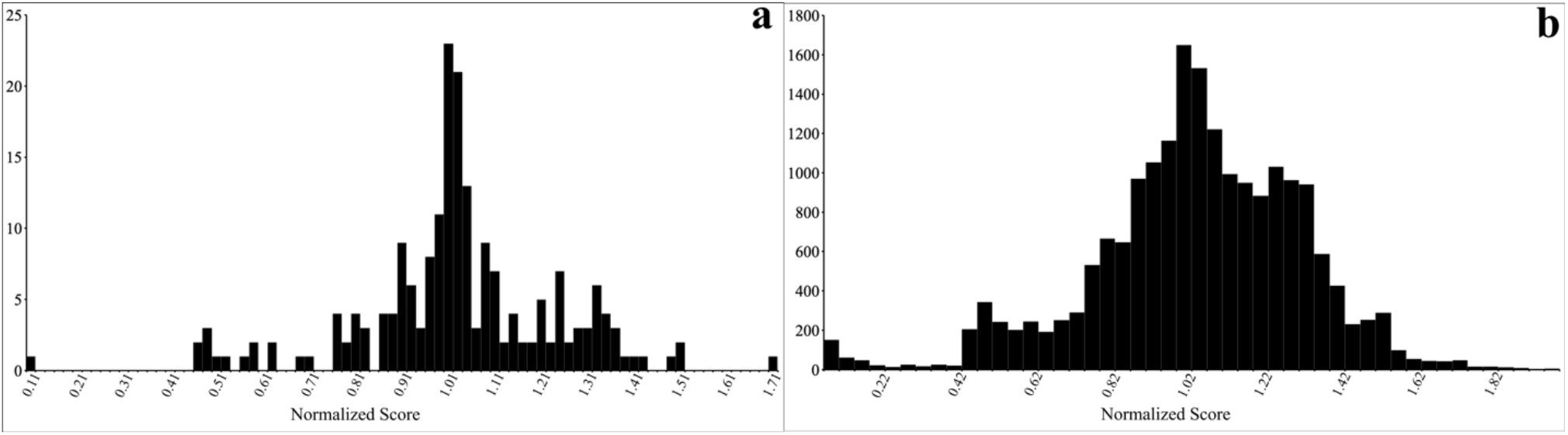
Histogram of network score after single perturbation (a) and double perturbation (b).

In Table 3 we list the perturbations that are consistent with experimental data and also result in a score lower than the score computed for the original network. The single-edge perturbations CycA-to-CycE: 0 to -1 and CycA-to-CycE: 0 to -1 cause large drops in the network score. The effect of both perturbations is to turn off the CycE activity after the cell cycle has entered the S phase. The interaction corresponding to the perturbation CycA-to-CycE: 0 to -1, namely, negative autoregulation of CycE, was indeed observed in Ref [20], where autophosphorylation of CycE-Cdk2 complex results in ubiquitin-mediated degradation of CycE. The interaction corresponding to the perturbation CycA-to-CycE: 0 to -1, namely negative regulation of CycE by CycA does not exist in databases. However, the known sequence of interactions CycA ⊣ Cdh1 ⊣ Skp2 ⊣ CycE results in the indirect negative regulation of CycE by CycA. Since Skp2 is not a part of Model B, we think the perturbation CycA-to-CycE: 0 to -1 tries to capture the indirect negative interaction with a direct one. Interestingly, the perturbation pRB-to-CycE: 0 to -1 is also present in our network for the Model C (Fig. 1c). Further, Model B has a lower graph score when the negative regulation of Cdh1 by CycE is removed from the network. This interaction is not found in databases and also not included in Model C (Fig. 1c).

**Table 3:** List of plausible perturbations resulting in a network score smaller than the original network.

| Perturbation | Normalized Graph Score | Steady state count | Largest attractor size | Correct | Incorrect | Did-not-start |
| --- | --- | --- | --- | --- | --- | --- |
| **OG Graph** | **1** | **100** | **283** | **58%** | **0%** | **42%** |
| Single Perturbation |  |  |  |  |  |  |
| CycE-to-CycE: 0 to -1 | 0.46 | 186 | 640 | 51% | 3% | 46% |
| CycA-to-CycE: 0 to -1 | 0.49 | 338 | 627 | 55% | 0% | 45% |
| pRB-to-CycE: 0 to -1 | 0.61 | 152 | 638 | 51% | 2% | 47% |
| CycE-to-Cdh1: -1 to 0 | 0.91 | 130 | 282 | 57% | 0% | 43% |
| Double Perturbation |  |  |  |  |  |  |
| CycE-to-CycE: 0 to -1<br>CycA-to-CycE: 0 to -1 | 0.45 | 126 | 642 | 50% | 4% | 46% |
| PPs-to-p21-p27: 1 to 0<br>Cdh1-to-Cdh1: 0 to 1 | 0.93 | 123 | 270 | 64% | 0% | 36% |

When both network edges CycA-to-CycE: 0 to -1 and CycA-to-CycE: 0 to -1 in Model B are simultaneously modified the graph score drops a little bit lower compared with the score results for corresponding single-edge perturbations. The graph score is also better when the activation of p27 by phosphatases PPs is removed and self-activation loop to Cdh1 is added. The last regulation is also present in Model C (Fig. 1c). Overall, almost all interaction modifications that give lower graph scores for Model B in our perturbation analysis are already present in Model C that we analyze next.

### Mammalian cell: Model C

Finally, we constructed a Boolean model using the ODE-based Goldbeter model of mammalian cell cycle [2]. The network of interaction is shown in Figure 1c. Compared to Model B in Figure 1b, this network has additional nodes Skp2, Wee1, and Cdc25. For this model we find that the dynamics of almost all the steady states converge to the attractor corresponding to G0 state listed below.

| CycD | CycE | CycA | CycB | E2F | Skp2 | Cdh1 | Cdc25 | pRB | p21-p27 | Cdc20 | Wee1 | PPs | Avg Count |
| --- | --- | --- | --- | --- | --- | --- | --- | --- | --- | --- | --- | --- | --- |
| 0 | 0 | 0 | 0 | 0 | 0 | 1 | 0 | 1 | 0 | 0 | 1 | 0 | 8192 |

The distribution of scores after single-edge and double-edge perturbations are shown in Fig. 4. The original network gives excellent results, and as a consequence, relatively few perturbations resulted in a lower score. The full list of network scores corresponding to single-edge and double-edge perturbations is given in the Supplementary Information File S3.

**Figure 4.**
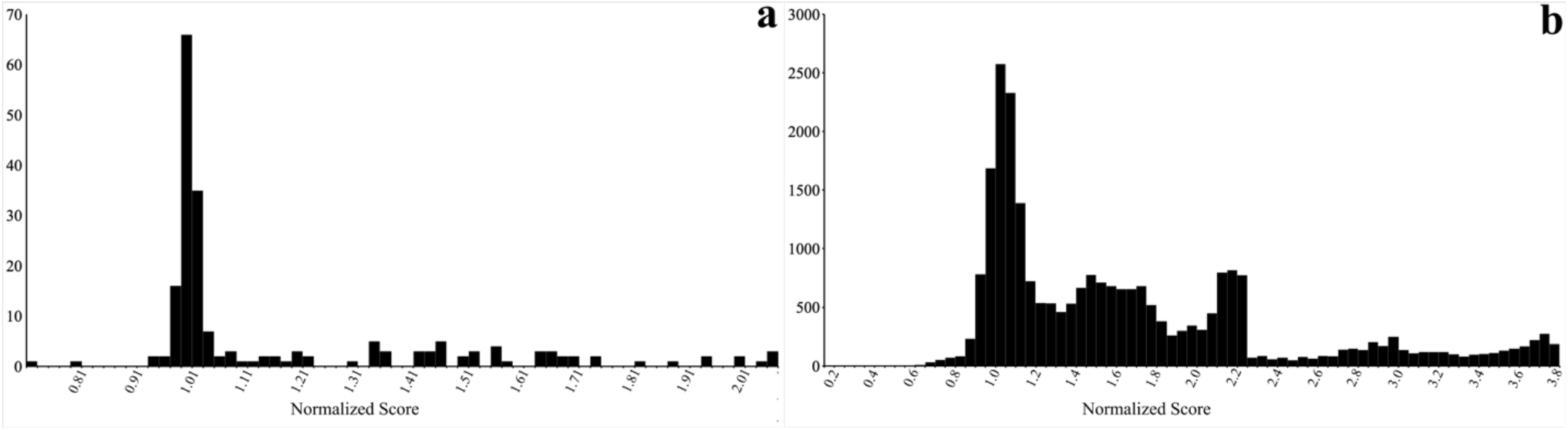
Histogram of network score after single perturbation (a) and double perturbation (b).

The perturbations that are consistent with experimental data and also result in a score lower than the score for the original network are listed in Table 4. The double-edge perturbation CycA-to-CycE: 0 to -1 and RB-to-RB: 1 to 0, resulted in the lowest graph score. Remarkably, the indirect negative regulation of CycE by CycA (CycA ⊣ E2F1 →CycE) is present in Model B. This double-edge perturbation improved the frequency of starting the cell cycle as well as the frequency of correct trajectories. Interestingly, the single perturbation RB-to-RB: 1 to 0 has a score of 13.8, but when combined with CycA-to-CycE: 0 to -1, results in a score of 0.85. The second lowest graph score was obtained by adding a negative interaction from phosphatases PPs on Skp2 and a negative regulation of p21 and p27 by Skp2. Both perturbations are found in databases. For example, dephosphorylation of Skp2 by the mitotic phosphatase Cdc14B promotes the degradation of Skp2 [21]. Also, Skp2 is required for ubiquitin-mediated degradation of p27 and p21 [22,23]. The double-edge perturbation Cdc25-to-CycE: 0 to 1 and Cdh1-to-Cdc25: 0 to -1 also improves the graph score. The up-regulation of CycE by Cdc25A has been confirmed in Ref [24]. It has been also observed that Cdc25A degradation is mediated by the anaphase-promoting complex (APC/C)(Cdh1) [25]. The final suggestion in the list of Table 4 is pRB-to-CycA: -1 to 0 and PPs-to-Wee1: 0 to 1 double-edge perturbation. The positive regulation of Wee1 by phosphatases agrees with studies reporting that phosphatase Cdc14A inhibits Wee1 degradation through dephosphorylation [26]. Also, the direct negative regulation of CycA by pRB does not appear in databases. Further, Model C has already a negative indirect influence of pRB on CycA (pRB ⊣ E2F1 →CycA). Thus, the direct inhibition of CycA by pRB is redundant.

**Table 4:**
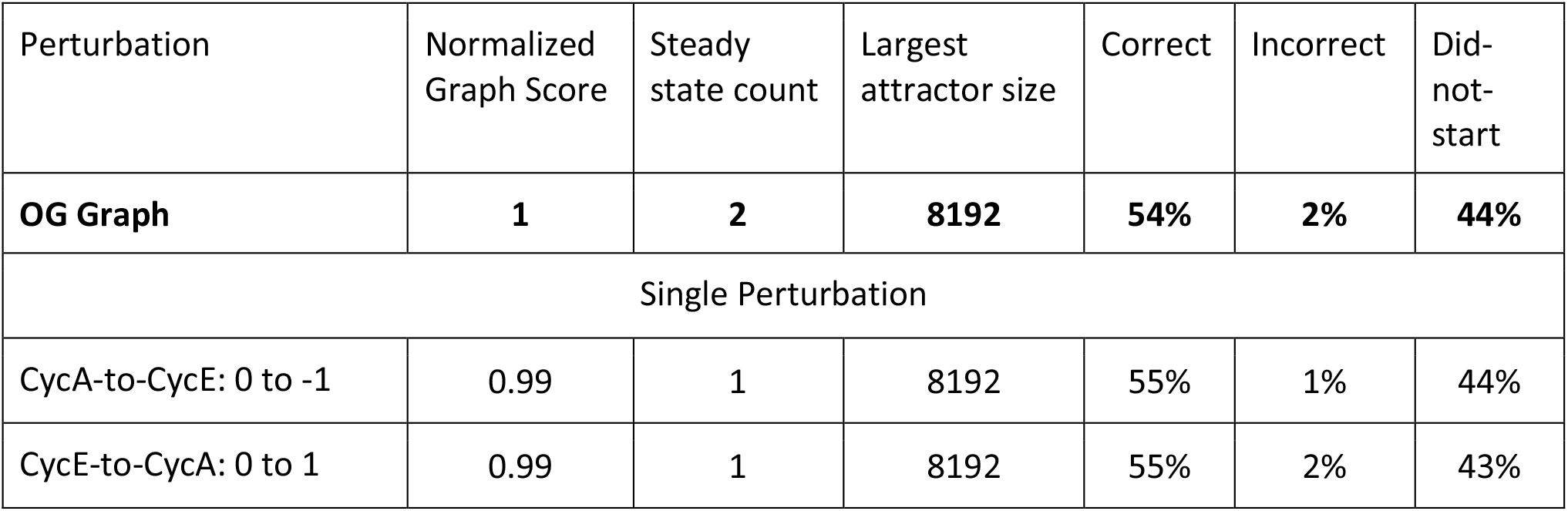
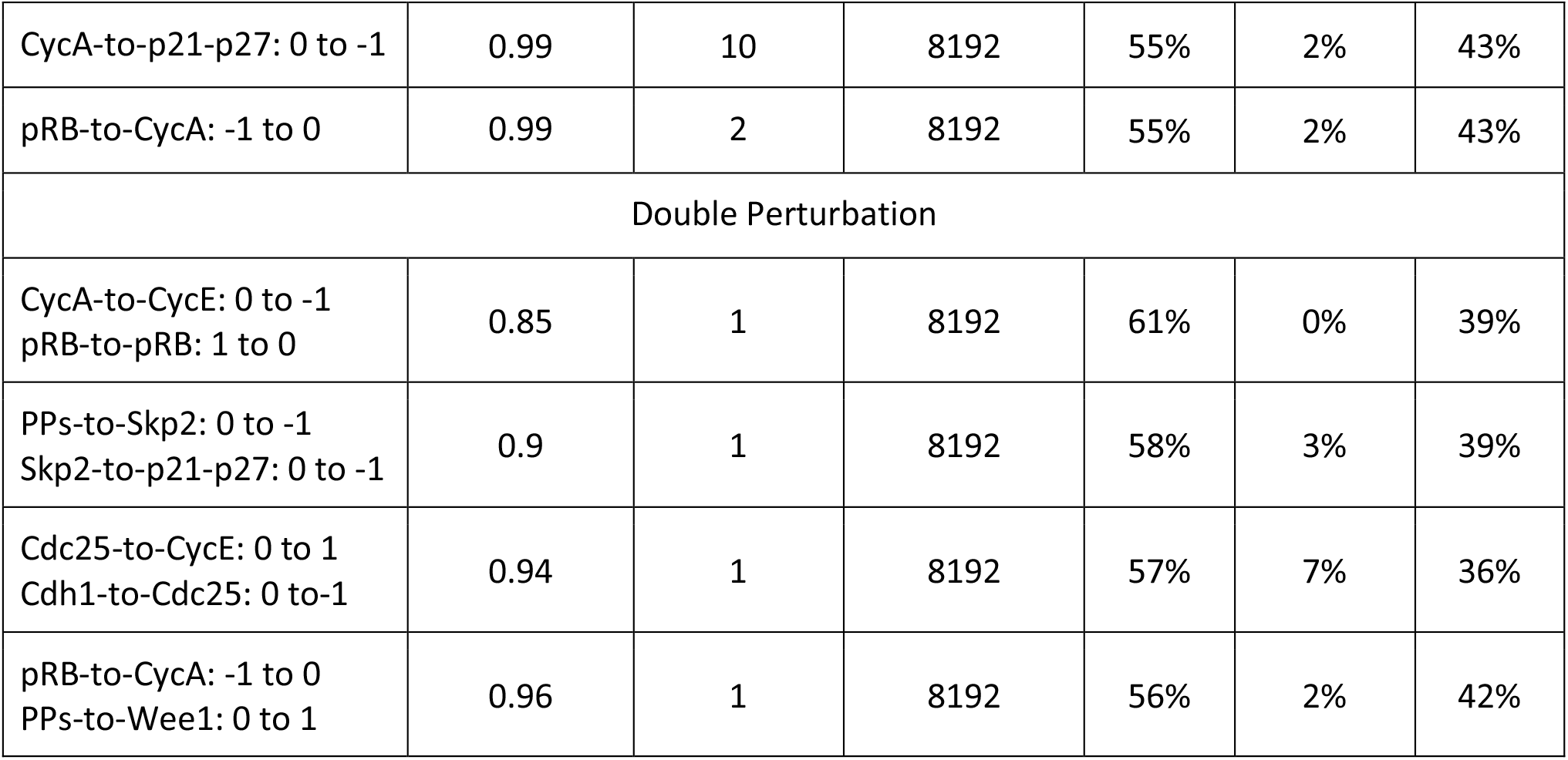
List of plausible perturbations resulting in a network score smaller than the original network.

## DISCUSSION

Understanding the complex dynamics of cell cycle is of both theoretical and practical interest. Lot of effort has been put into developing ODE-based and Boolean dynamical models of cell cycle, that are consistent with experimental data. The main challenge in developing such dynamical models is the lack of a systematic approach in putting together the model, and later incorporating new genes and proteins in the model. The absence of such a structured approach is apparent among existing cell cycle models, which exhibit differences in their sets of variables and the interaction networks connecting them.

In this manuscript we developed a semi-automated method to construct and improve Boolean models of cell cycle. The main steps in our approach are the following: (a) manually construct an initial Boolean model using existing Boolean or ODE models of cell cycle (b) define a network score to quantify the consistency of network dynamics with that of cell cycle, (c) using single and double perturbations of the network, find interactions that result in a lower score, (d) compare the proposed interactions in (e) with databases to find the ones that are consistent with experimental data, and finally (e) manually go through the interaction in (d) to ascertain their validity. Couple of important points: first, while going through step (e) we found that the protein-protein interactions listed in the databases are not entirely consistent with the experimental data available in the literature. Hence a manual screening of the interactions was necessary. And second, the steps described above can be used to add new protein to the network and extend the models.

We first applied our perturbation method to the well-known model of cell cycle in budding yeast. Through double perturbation analysis we found interactions that significantly improved the size of the largest attractor as well as the frequency of correct transitions through the cell cycle trajectory. Interestingly, these interactions (Cln1,2-to-MBF: 0 to 1, Clb5,6-to-Mcm1: 1 to 0) are not present in any database, but since they appeared naturally from our analysis, searching for them in the literature became much easier. We then applied the same method to two different models of cell cycle in mammalian cells – the Tyson-Novak and Goldbeter model. In both cases we found perturbations that improved the network score.

The number of double perturbations increases as N^4^ with the number of nodes in the network. Combining it with the fact that for each network perturbation multiple simulation runs are needed to compute the statistical properties of the dynamics, the computation time increases significantly with the number of nodes. However, we found that retaining only the feasible single perturbations for the double perturbation analysis, significantly reduced the computational burden. For example, for Model C, about 80% of all single perturbations were classified ‘false’ after database search, and retaining only the ‘true’ single perturbations for the double perturbation analysis reduced the search to only about 4% of all possible double perturbations.

## METHODS

### Cell-Cycle Models and Update Scheme

The Boolean models of cell cycle contain nodes and directed edges. The nodes represent proteins and the directed edges correspond to the interactions between them. The edges are of two different types, inhibitors (−1 weight) and activators (+1 weight). The edges can also be self-loops, i.e. directed edges starting and ending at the same node.

We use the asynchronous update scheme to calculate the network dynamics. At each time step, a node is selected randomly and its value is updated using the following rule

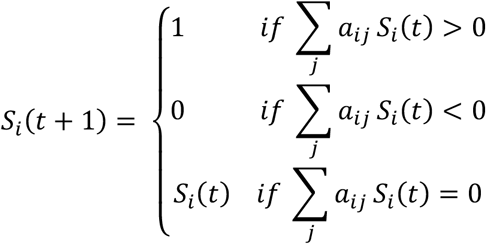

Here *S*_*i*_(*t*)= 0 *or* 1 corresponds to the value of node *i* at time *t*, and *a*_*ij*_ are the weights of the edges joining node *i* to node *j*. We chose *a*_*ij*_ = 1 for activation and *a*_*ij*_ = −1 for deactivation. Since an asynchronous update scheme results in stochastic trajectories, we simulated multiple trajectories for each initial condition to obtain the statistical details of the model dynamics. We follow the same update scheme throughout the publication, except where explicitly mentioned.

### Sequence

For each model, we fix a protein whose activation is considered as the start of the cell cycle. We call the activation of this protein as the point-of-no-return. If in a simulation the protein is not activated and the trajectory reaches the steady state corresponding to G0, we classify the trajectory as ‘did-not-start’. If the dynamics passes the point-of-no-return, we check for a predefined order of protein activation to classify the dynamics as correct or incorrect. If the dynamics progresses via the predefined sequence of states, we classify the trajectory as ‘correct’, and otherwise ‘incorrect’. The point of no return, G1 states, and the correct sequence for different models are given below.

#### Model A (Budding Yeast Model)

In the network shown in Figure 1a, the cell size acts as the start signal. In the absence of this signal the cell remains in G0 phase (a steady state of the model), with the inhibitors of the cell cycle Cdh1 and Sic1 being active. In the presence of the signal the cell enters the G1 phase where Cln3 is activated. At this point the cell cycle can move forward to the S phase by activating Clb5,6 or go back to the G0 phase (we refer to this as the did-not-start case). Upon Clb5,6 activation the cell is committed to go through the cell cycle. The cycle is characterized by the sequential activation of Cln3, Cln1,2, Clb5,6, and Clb1,2.

Point of no return: Clb5,6 activation

G1 states: [Cln3, Cdh1, Sic1] = 1 and [Cdc20,14, Clb5,6, Cln1,2, Clb1,2, Mcm1, SFF] = 0.

Correct sequence: [Clb5,6=1, Clb1,2=0, Cdc20,14=0] → [Clb1,2=1, Cdc20,14=0] → [Cdc20,14=1]

#### Model B (Modified Tyson-Novak Mammal Model)

Point of no return: CycE activation

G1 states: [CycD, RB, P27, Cdh1] = 1 and [CycE, CycA, CycB, Cdc20] = 0.

Correct sequence: [E2F=1, CycE=0, CycA=0, CycB=0, Cdc20=0] → [CycE=1, CycA=0, CycB=0, Cdc20=0] → [CycA=1, CycB=0, Cdc20=0] → [CycB=1, Cdc20=0] → [Cdc20=1]

#### Model C (Modified Goldbeter Mammal Model)

Point of no return: CycE activation

G1 states: [CycD, RB, Wee1] = 1 and [CycE, CycA, CycB, E2F, Cdc20] = 0.

Correct sequence: [E2F=1, CycE=0, CycA=0, CycB=0, Cdc20=0] → [CycE=1, CycA=0, CycB=0, Cdc20=0] → [CycA=1, CycB=0, Cdc20=0] → [CycB=1, Cdc20=0] → [Cdc20=1]

### Perturbations and Scoring

Single perturbations in the network were introduced by changing an edge value using the following possibilities: 0 → (1 and 1), 1 → (1 and 0), 1 → (1 and 0). We analyzed all possible perturbation to all edges (including self-loops), resulting in a total of 2*N*^2^ single perturbations. Double perturbations were introduced by applying single perturbation to two edges at a time.

To rank the performance of perturbed networks, we defined a scoring scheme that incorporates penalty if any initial condition did not reach G0 after starting the cell cycle, or if the sequence of activation/deactivation events was not consistent with the sequence known for the cell cycle (initial states corresponding to G1 states only), or if the cell cycle goes to G0 without starting the cell cycle (we call it did-not-start case).

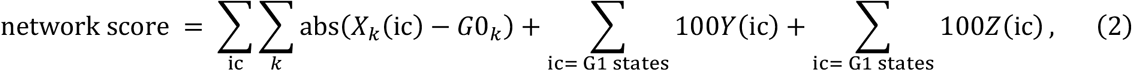

where *X*_*k*_(*ic*)is the *k*^*th*^ component of the steady state corresponding to in the initial condition *ic*, and *G*0_*k*_is the *k*^*th*^component of the quiescent G0 state. The quantity *Y* is for sequence-related penalty; it takes the value 0 if the sequence is correct and 1 if the sequence is incorrect. The quantity *Z* is the penalty associated with the did-not-start case. **Z** takes the value 0 if the cell cycle starts and 1 if the cell goes back to G0 instead of going to the S phase. The sequence and did-not-start related penalties were introduced only for G1 states.

## ACKNOWLEDGEMENT

The work was supported by the Academy of Data Science Discovery Fund awarded to AB and PK.

## AUTHOR CONTRIBUTIONS

AB and PK conceptualized the project. AR wrote and implemented Python scripts to do single-edge and double-edge perturbations analysis of Boolean networks. AM wrote scripts to compare new interactions suggested by simulations with those available on protein-protein interaction databases. AB, PK and AR prepared the manuscript.

## DATA AVAILABILITY

Python codes used for the simulations and for generating figure panels can be accessed on GitHub: https://github.com/asif256000/boolean_cellcycle_analysis/releases/tag/v1.0

## Notes

https://github.com/asif256000/boolean_cellcycle_analysis/releases/tag/v1.0

### Competing Interest Statement

The authors have declared no competing interest.

