## Supplementary Information for "A Perturbation Approach for Refining Boolean Models of Cell Cycle Regulation"

**Supplementary File S1**

This Excel file contains statistical data related to the single-edge (sheet 1) and double-edge (sheet 2) perturbation analysis of Model A (cell cycle in budding yeast). The data we calculated after performing 500 iterations per single-edge perturbations and 20 iterations per double perturbation. Sheet 3 shows the final states of the model after and their corresponding sizes. The calculation of final states is described in the execution flow below.

**Supplementary File S2**

This Excel file contains statistical data related to the single-edge (sheet 1) and double-edge (sheet 2) perturbation analysis of Model B (cell cycle in mammalian cell based on the Tyson-Novak model). The data we calculated after performing 500 iterations per single-edge perturbations and 20 iterations per double perturbation. Sheet 3 shows the final states of the model after and their corresponding sizes. The calculation of final states is described in the execution flow below.

**Supplementary File S3**

This Excel file contains statistical data related to the single-edge (sheet 1) and double-edge (sheet 2) perturbation analysis of Model C (cell cycle in mammalian cell based on the Goldbeter model). The data we calculated after performing 500 iterations per single-edge perturbations and 20 iterations per double perturbation. Sheet 3 shows the final states of the model after and their corresponding sizes. The calculation of final states is described in the execution flow below.

**Description of columns in the Supplementary Files**

**Graph Modification ID:** This column signifies the perturbation to the original model, and the original model is denoted by 'Original Model' in this column. For other rows, the column follows the pattern "*<source node>-to-<target node> -> <from value>to<to value>*", and for multiple perturbations the patterns are joined by " | " to signify separate perturbation. For example, "CycA-to-CycE -> 0 to1 | CycA-to-P21 -> 0 to-1" means that for this particular perturbation we changed the weight of the edge going from Cyclin A (CycA) to Cyclin E (CycE) from 0 to 1 AND the weight of the edge going from Cyclin A (CycA) to P21-P27 (P21) from 0 to -1.

**Exists in Database:** This column can only be found for mammal models (Model B and Model C). We checked the validity of the perturbations generated by simulations by comparing the interaction against the protein-protein interaction database SIGNOR (https://signor.uniroma2.it). a. For the case of a single perturbation, we checked if the corresponding entity pair (node1, node2, interaction) exists in the database. If they do, we mark the perturbation as 'True', and otherwise 'False'. In cases of self-loops (edge going from a node to itself), we do not have any data in the database. In such cases we mark the column as 'NA'. Also, if the perturbation is removal of an edge (perturbation from 1/ -1 to 0) and we don't find any interaction between those two nodes in the database, then we mark the column as 'True' as it implies there is no interaction between those two nodes. In case of double perturbation, we mark this column as 'True' only if both the perturbations align with the results from the database. If one of the perturbations in double perturbation is self-loop, we ignore the self-loop and mark this column based on the result from the other perturbation.

**Database Context:** This column we provide some context for the perturbations for which we find a match in the database. The format of the 'DB Context' column is '*[<db_source_node> -> <db_target_node> -> <interaction_type> -> <pubmed_id(s)>]*'. Here 'db_source_node' is the node name found in the database that matches with the source node in the perturbation, 'db_target_node' is the node name found the database that matches with the target node in the perturbation, 'interaction_type' can be either 1 or -1 based on their interaction type (activation/repression). The 'pubmed_id(s)' are the pubmed_id(s) that support the interaction given in the database. In case of double perturbation, wherever possible, we write two such 'DB Context' in the same row.

**Absolute Graph Score:** We calculate the 'Absolute Graph Score' based on the formula given in 'Perturbations and Scoring' section under 'Methods'. The exact formula to calculate the score is given in the 'Methods' section.

**Normalized Graph Score:** To calculate the 'Normalized Graph Score', we divide the 'Absolute Graph Score' of each perturbation with the absolute score of the 'Original Model' for that particular model. For example, if we have the 'Absolute Graph Score' for some perturbation as 1200, and the 'Absolute Graph Score' for the 'Original Model' of the same model is 800, then we can calculate the 'Normalized Graph Score' for the particular perturbation as (1200/ 800) = 1.5. By definition the 'Normalized Graph Score' for the 'Original Model' for each model is always 1.

**Steady State Count:** This is the average number of final state(s) for each model after executing them for a particular number of iterations. Because of the stochastic update method and finite number of iterations, in some simulation runs the dynamics does not reach the steady state. Therefore the 'steady state count' can be slightly larger than the number of actual steady states of the model.

**Largest Attractor Size:** This is the average size of the largest final state. For example, a graph with 10 nodes has $2^{10} = 1024$ initial states, each of which reach some final state at the end of the simulation run. If there are 3 'Steady States' for this cycle with the distribution being 1000, 18, 6, then we say that the 'Largest Attractor Size' for this model to be 1000.
**Correct(%), Incorrect(%), Did not Start(%)**: These columns denote the percentage of trajectories (with initial condition limited to G1 states) that go via the correct cell cycle sequence, incorrect cell cycle sequence, or do-not-start the cell cycle at all, respectively. More details of the process to distinguish them are discussed in detail in the 'Sequence' section under 'Methods' header.

Note: As the database we used to verify the perturbations contain only interactions for mammals, we do not have that data for the yeast model. Hence the supplementary materials for the 'Model A' does not contain the columns 'Exists in DB' and 'DB Context'.

**Most Frequent Steady State(s):** This is the final state corresponding to the 'Largest Attractor Size' for each graph.

**Execution Flow of the cell cycle:**

We follow a particular method to execute the simulations for each model and gather statistical data from them. For each 'Model' with 'n' nodes, there can be a maximum of $2^n$ initial conditions (IC1, IC2, … , IC$2^n$), which are simulated to generate a score for that execution using the formula given in the 'Perturbation and Scoring' section. We repeat the steps multiple times, gathering scores and other relevant data for every complete execution. The number of execution (x) to gather this statistical data varies based on the time taken to execute a complete cycle. After gathering data from all such 'x' executions, we perform statistical operations on the data to make them presentable.

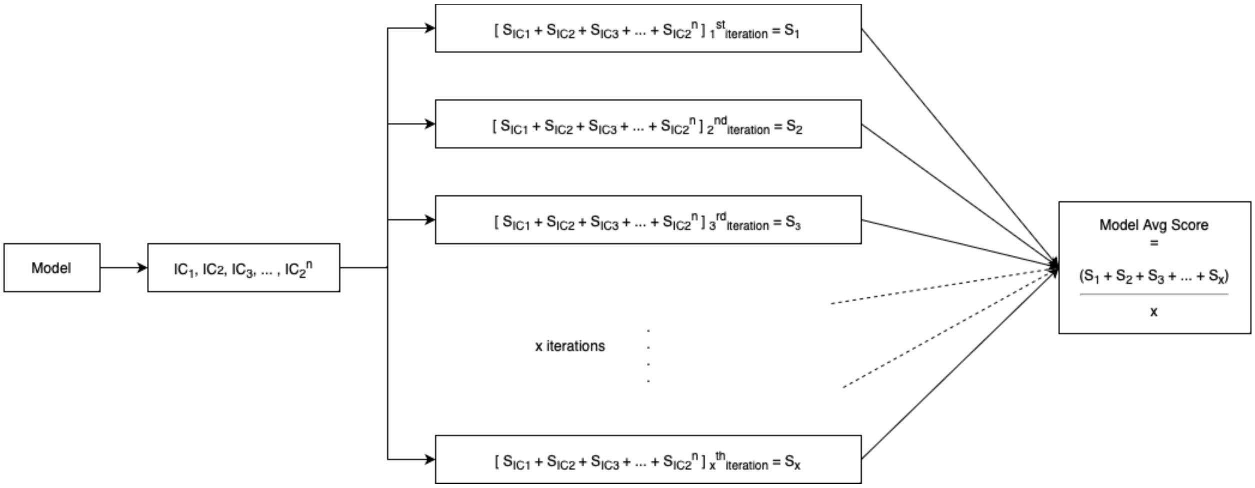


Figure 2: Execution Flow for a single model

Due to computational limitations, we do not execute every cell cycle till it reaches a true steady state. Rather we impose an upper bound on the number of updates each cell cycle can have under different conditions. For example, when we are calculating the performance of all perturbations we limit each cell cycle to a maximum of $(100 * Number\ of\ nodes)$ updates. That means, for each initial state $(IC1, IC2 \ldots IC2^n)$, we make updates to the cell cycle for $(100 * \ Number\ of\ nodes)$ times and the final state achieved after these updates are considered the steady states for the initial states. Again when we are looking for the true steady states, we have to let the cell cycle run for a lot more steps, sometimes around $(500 * Number\ of\ nodes)$ times. Even after all these updates, we have to manually take another look at the final states and individually analyze them to verify if they are true steady states or not.

**Steady States:** As the update method is stochastic in nature, due to finite number of iterations, the number of steady states can differ between two simulation runs. We put an upper limit on the number of iterations to narrow down the final steady states, and then checked for the true steady states by choosing this set as the initial condition and increasing the number of iterations.

**Cycle Detection:** The model can also detect whether there is a simple cycle at the end of a cell cycle. For any initial state, we run the cell cycle for at least 100*n times, where n is the number of nodes in each model. After the cell cycle is iterated for the number of given iterations, if sequences of states repeats in the same order multiple time, we conclude that the cell cycle has gone into a cycle and define the final state as "C" for every node, e.g. (CycD: C, CycE: C, … ). Even though we tried detecting most of the cycles, there are more convoluted cycles visible in the first mammal model that are very difficult to identify.
